# The elusive resistome: a global comparison reveals large discrepancies among detection pipelines

**DOI:** 10.64898/2026.05.11.724158

**Authors:** Juan S. Inda-Díaz, Faith Adegoke, Ulrike Löber, Víctor Hugo Jarquín-Díaz, Yiqian Duan, Johan Bengtsson-Palme, Svetlana Ugarcina Perovic, Luis Pedro Coelho

**Author notes:** **Corresponding authors** Correspondence to Luis Pedro Coelho and Juan S. Inda-Díaz.

## Abstract

Identifying antibiotic resistance genes (ARGs) from metagenomic data is critical for studying antimicrobial resistance across microbial communities and pathogens. However, there is no standardized methodology for ARG annotation. Here, we compare ten commonly used ARG detection pipelines by analysing over 270 million prokaryotic genes from the Global Microbial Gene Catalogue across 13 distinct habitats. We observed up to a 45-fold difference in the number of reported ARGs, with a mean Jaccard index of only 16% between pipelines. Pipeline selection profoundly impacted downstream biological interpretations, with drastic changes to estimates of ARG relative abundance and richness, to the characterization of pan- and core-resistomes, and to the class-level composition of the inferred resistome. ARG detection pipelines make different, defensible trade-offs, and no single approach should be treated as authoritative. Therefore, users should justify and communicate choices carefully, as our analyses show that, taken uncritically, the same data can support conflicting biological and ecological interpretations.

## Introduction

Antimicrobial resistance (AMR) represents an escalating global health challenge, causing over one million deaths in 2021 and, if no actions are taken, projected to cause ten million deaths annually by 2050^1,2^. AMR is primarily associated with antibiotic resistance genes (ARGs), genes that encode a range of functions that protect cells against antimicrobials^3,4^. Since the introduction of human-made antibiotics, the mobilization of ARGs from their natural sources into clinically relevant bacteria and mobile genetic elements has accelerated^5,6^. Despite their clinical importance, the global distribution and evolutionary origins of most ARGs remain poorly characterized^7–9^. Moreover, as the global threat of AMR grows, there is yet no standardized methodology for routine, genomic-based surveillance^10^, which is recognized as an obstacle to the implementation of genomics in public health efforts to mitigate AMR^11^.

Metagenomics is commonly used to characterize the presence of ARGs in host-associated and environmental samples^12^. The ARGs found in a microbial community, commonly known as the resistome, can be characterized by their richness, the number of unique ARGs, and their relative abundance, the fraction of the metagenome that encodes ARGs^13^. Across a group of samples, the pan-resistome refers to the ARGs found in any sample, while the core-resistome refers to those ARGs found in most samples in the group; the exact definition depends on the study^14–18^. Additionally, ARGs can be grouped into classes based on the antibiotics to which they confer resistance, their mechanisms of action, or their phylogeny^19–21^. This allows quantification of the resistome at the class level, thereby aiding interpretability^22^.

Numerous tools exist for identifying ARGs from genomic or metagenomic sequencing data, differing not only in their detection approaches, including sequence alignment, deep learning, and profile Hidden Markov Models, but also in reference databases. Consequently, these tools implement different, and sometimes unclear, criteria for identifying ARGs, and consensus remains lacking on tool selection and their application to specific research aims, including resistome quantification. Among these approaches, alignment-based methods include the Resistance Gene Identifier (RGI)^23–25^, ABRicate^26^, and ResFinder^27,28^. DeepARG expands on these tools by implementing an approach that, after pre-detection by alignment, uses deep learning to annotate genes into resistance classes^29^. Other tools employ profile Hidden Markov Models (HMMs) to capture broader protein families^30^; for example, fARGene^31^ utilizes optimized HMMs that balance sensitivity and specificity using true positive and true negative resistance gene references to detect several major ARG classes^31–36^, while AMRFinderPlus uses both sequence alignments and HMMs^37^.

The output of these tools is fundamentally tied to reference databases. These include the Comprehensive Antibiotic Resistance Database (CARD)^23,38,39^, organized under the Antibiotic Resistance Ontology (ARO) framework^40^; NCBI’s reference collection, which catalogs genes and point mutations alongside HMM profiles associated with resistance^37^; and ResFinder, which focuses on acquired ARGs^27,28^. Other ARG databases, such as ARG-ANNOT (not actively updated)^41^, further expand this landscape, and meta-databases like MEGARes^42^ aim for comprehensiveness by integrating sequences from CARD, ResFinder, NCBI, and the BacMet Antibacterial Biocide and Metal Resistance Genes Database^43^. Some tools are designed around a single reference database, whereas others support multiple databases; all allow user-defined parameter settings, creating an enormous number of potential approaches to identify ARGs.

In this study, we evaluated how selection of the ARG detection pipeline influences our view of the resistome–from relative abundance and richness to the characterization of pan- and core-resistomes across host-associated and external microbial habitats. Here, we define a “pipeline” as a specific combination of tool, database (where applicable), and parameter settings. We leveraged the Global Microbial Gene Catalogue (GMGCv1)^44^–a set of over 270 million non-redundant representative sequences (unigenes) derived by clustering 2.3 billion prokaryotic sequences at 95% nucleotide identity–for analysis using ten different ARG detection pipelines. We assessed the variability in resistome estimates across pipelines and identified methodological effects on the detection of specific ARG classes. Our findings demonstrate how pipeline selection can lead to conflicting biological interpretations.

## Results

We analyzed 278,788,551 microbial unigenes from GMGCv1 with ten different ARG detection pipelines: RGI, DeepARG, fARGene, ResFinder, and AMRFinderPlus with default databases and models, and ABRicate with five databases. We found 178,107 unigenes (0.063%) reported as ARG by at least one pipeline. We explored the presence of these unigenes in 11,538 metagenomic samples across 13 distinct habitats, including human-associated (gut, skin, nose, oral, and vaginal), non-human host-associated (pig, mouse, dog, and cat), and external habitats (wastewater, marine, freshwater, and soil; Supplementary Table 1). We quantified ARG relative abundance and richness in metagenomic samples as well as pan- and core-resistome profiles across habitats, using the output of each pipeline.

### Variation in the size of the resistome

The total number of reported ARGs varied drastically across pipelines; for instance, ABRicate using the ResFinder database (ABRicate-ResFinder) reported only 2,213 ARGs, whereas DeepARG reported 100,075—a 45-fold difference (Fig. 2a). The mean overlap between tools, measured by the Jaccard index, was as low as 16% (Fig. 2c). These differences were not solely driven by the underlying databases but also by pipeline-specific detection criteria. RGI, which utilizes CARD, reported more ARGs than ABRicate using CARD (ABRicate-CARD), with the latter capturing only 8.2% of RGI’s predictions. Conversely, RGI only reported 53.8% of the ARGs flagged by ABRicate-CARD. Similar discordance was observed with the NCBI and ResFinder databases: ABRicate using NCBI (ABRicate-NCBI) and ABRicate-ResFinder captured only 18.5% and 80.0% of the ARGs reported by AMRFinderPlus and ResFinder, respectively.

**Fig. 1:**
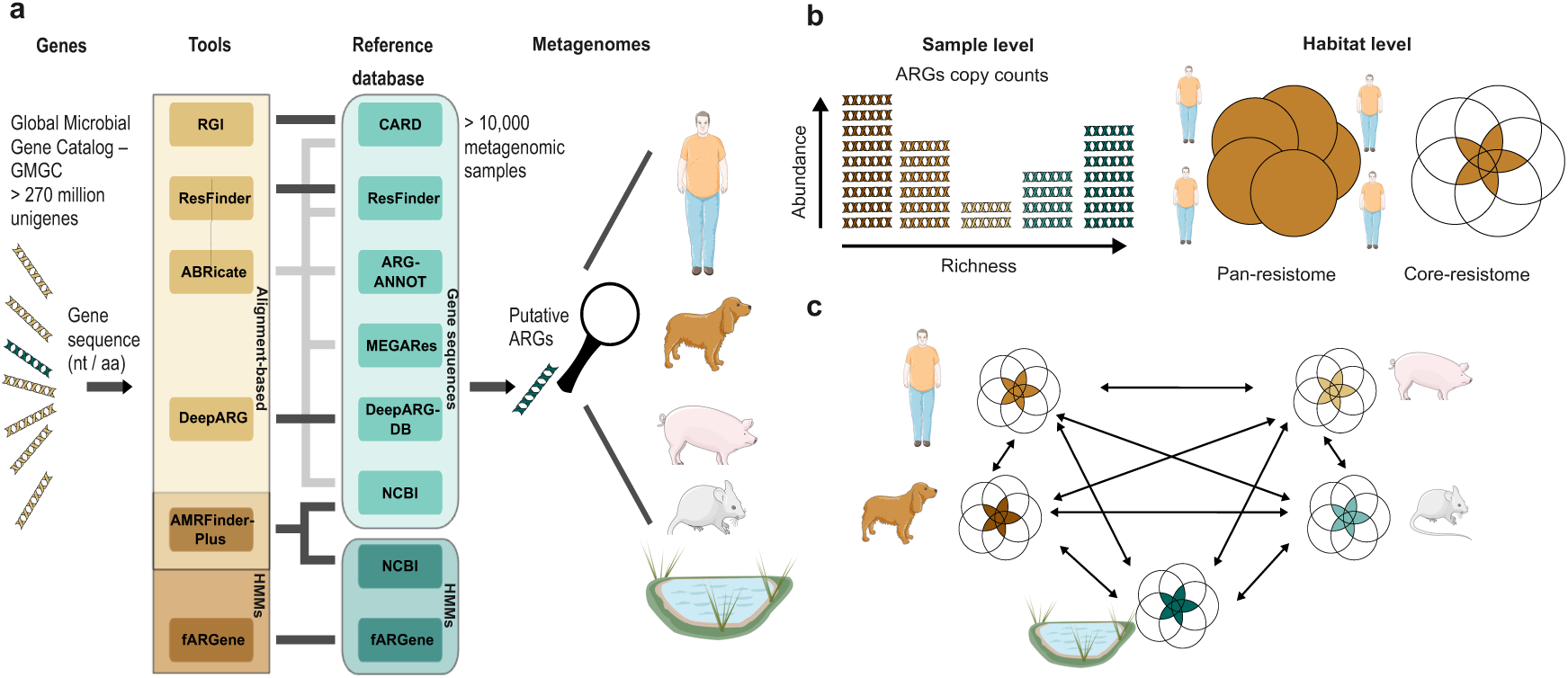
**Conceptual framework for resistome analysis**. **(a)** The GMGCv1 dataset is processed through multiple ARG detection pipelines to generate a comprehensive list of predicted ARGs. **(b)** Resistome metrics, including abundance, richness, and pan- and core-resistome sizes, are estimated from unigene profiles and their respective copy numbers across metagenomic samples and habitats. **(c)** Pan- and core-resistome analyses are employed to assess the intra- and inter-habitat dissemination of ARGs.

**Fig. 2:**
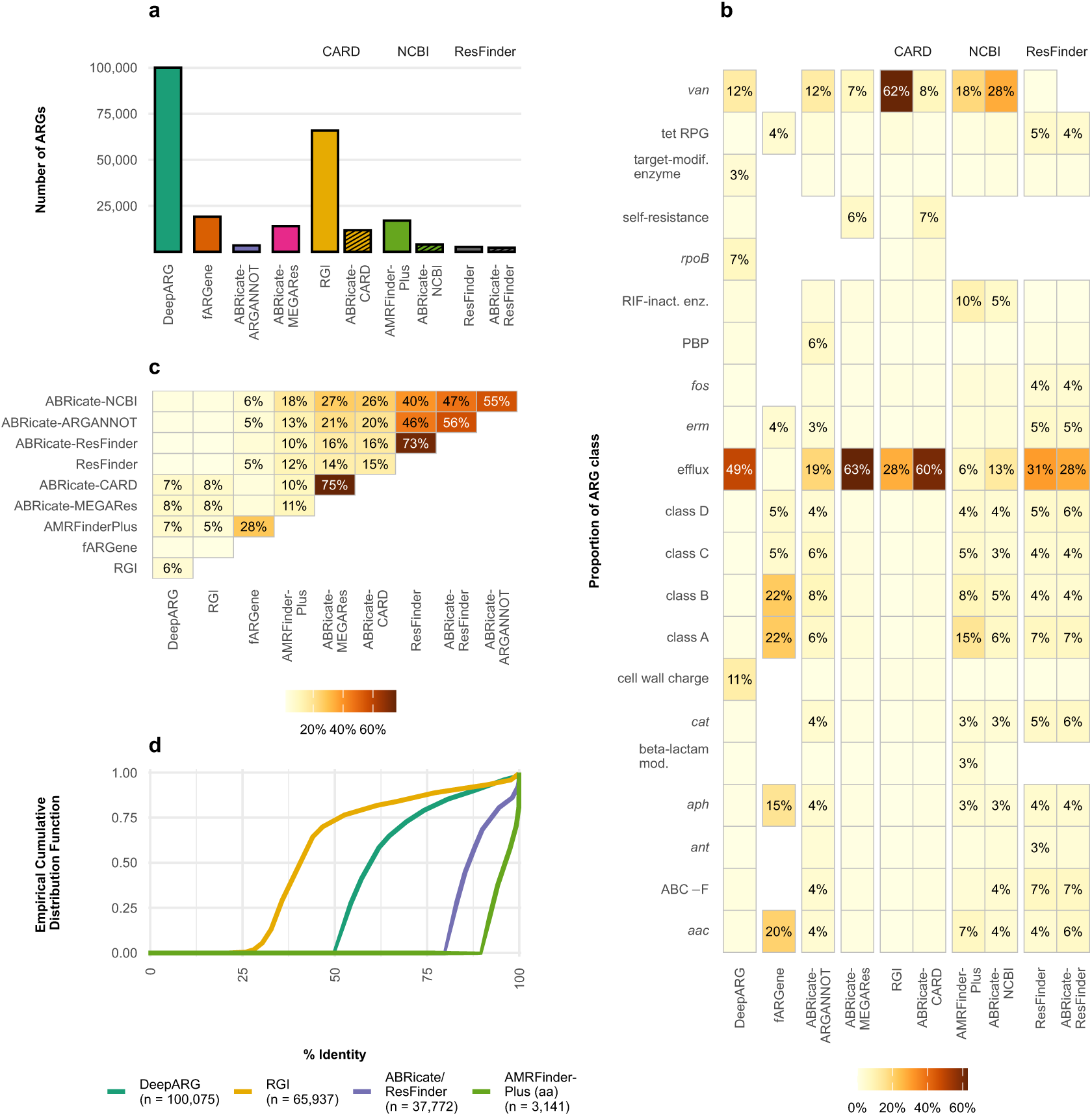
Differences in ARG counts and functional classification across detection pipelines. **(a)** Total number of ARGs reported by each pipeline. **(b)** Proportion of ARG classes reported by each pipeline; only classes representing ≥ 5% of the total within at least one pipeline are visualized. Missing cells indicate gene classes with no hits or not reported by the pipeline. **(c)** Pair-wise Jaccard index using the set of genes reported as ARG by each pipeline. **(d)** Empirical Cumulative Distribution Function of the sequence identity levels for ARGs reported via alignment-based methods. For RGI and DeepARG, 75% of reported ARGs had sequence identities below 51% and 71%, respectively. For AMRFinderPlus, only the subset of ARGs reported through protein alignment (14% of its total predictions) is included. Predictions from fARGene and the HMM-based component of AMRFinderPlus are excluded from this panel, as HMM-based detection does not provide a direct sequence identity score comparable to alignment. The groups “class A”, “class B”, “class C”, and “class D” represent 𝛽-lactamases, while “tet RPG” represents tetracycline ribosomal protection genes.

The majority of all reported ARGs were concentrated in two gene classes: efflux pumps (𝑛 = 63,964, 36%) and *van* genes, genes associated with a glycopeptide resistance cluster, (𝑛 = 52,767, 30%). These classes accounted for 92%, 87%, 75%, and 60% of the ARGs reported by RGI, ABRicate using MEGARes (ABRicate-MEGARes), ABRicate-CARD, and DeepARG, respectively, but less than 32% of those reported by ResFinder, AMRFinderPlus, and fARGene. Despite the significant difficulties in distinguishing between resistance-conferring pumps and those involved in general physiological transport^25^, efflux pumps comprised a substantial proportion of all reported ARGs. Notably, 23,784 (58%) of the efflux pumps identified by DeepARG were classified by the tool as “multidrug”–a category highlighted by the authors of the tool as an important technical challenge requiring manual curation^29^–and an additional 5,290 (13%) were labeled as “unclassified”. Similarly, glycopeptide resistance genes may be reported based on their relatedness to genes within the *van* gene cluster. However, distant homologs of these genes may instead be involved in cell-wall metabolism, regulation, or peptidoglycan biosynthesis, without necessarily conferring a vancomycin-resistance phenotype^45^. In contrast, 𝛽-lactamases constituted a substantial proportion of the ARGs identified by fARGene (53%), AMRFinderPlus (32%), and ResFinder (20%). In addition, aminoglycoside resistance genes, predominantly phosphotransferases (*aph*) and acetyltransferases (*aac*), accounted for 35% of the total ARGs reported by fARGene (Fig. 2b).

In general, alignment-based methods assume that gene function can be inferred from sequence similarity and rely on thresholds, such as sequence identity or bit-scores, to classify hits. A substantial majority of the ARGs reported by DeepARG (𝑛 = 73,649; 74%) and RGI (𝑛 = 56,290; 85%) exhibited less than 70% amino acid identity to their respective reference sequences (Fig. 2d). Crucially, 90% of these low-identity predictions were unique to the specific pipeline that reported them. In the case of RGI, we considered only “Perfect” and “Strict” matches, defined as exact matches and those passing curated similarity thresholds, respectively^39^. Nonetheless, 48,554 of the ARGs (74% of its total) exhibited <50% identity to the reference. These low-identity hits were predominantly concentrated in the *van* ( 𝑛 = 36,726 ), efflux pump ( 𝑛 = 10,533 ), and tetracycline ribosomal protection genes (RPGs; 𝑛 = 791) classes. Although lower identity thresholds facilitate the discovery of divergent or novel resistance genes, they also increase the risk of false-positive calls^46^. In fact, permissive thresholds did not result in more comprehensive ARG sets–neither encompassing those identified by more conservative pipelines nor improving sensitivity to mobile, and thus potentially clinically relevant genes, as DeepARG and RGI reported only 65% and 49% of ResFinder ARGs, respectively.

In some cases, antibiotic resistance is conferred not by the presence of a gene but by specific mutations in housekeeping genes^47^. For example, point mutations in the universal bacterial gene *rpoB*, which encodes the RNA polymerase-𝛽-subunit, can confer resistance to rifampicin^48–50^. Pipelines that annotate *rpoB* variants as ARGs solely based on sequence identity thresholds, without strict filtering for validated resistance-conferring mutations, are likely to overestimate the resistome. Notably, DeepARG reported 6,808 *rpoB* genes, all classified by the tool as “multidrug”, with 96% showing <80% amino acid identity to the reference gene, and without obvious filtering for mutations associated with resistance. Given that *rpoB* can serve as a proxy for genome count in metagenomic data^51^ and as a taxonomic marker^52^, the large number of *rpoB* variants reported by DeepARG likely reflects bacterial diversity rather than antibiotic resistance function, thereby inflating resistome estimates.

### Variation in the relative abundance of ARGs across pipelines

The relative abundance of ARGs was significantly higher across all habitats for DeepARG and RGI than for other pipelines. Their median relative abundance was at least two-fold higher in host-associated (excluding the human vaginal habitat) and wastewater habitats and at least five-fold higher in freshwater and marine samples compared to other pipelines (Fig. 3a; Extended Data Fig. 1). The largest differences occurred in the marine habitat, where ResFinder and ABRicate (across all databases) reported negligible resistance levels. Conversely, RGI estimated marine resistance levels exceeding those of soil—a result unique to this pipeline. This pattern was driven almost exclusively by efflux pumps and *van* genes, which accounted for a median of 99.8% of the total ARG relative abundance across marine samples reported by RGI (Extended Data Figs. 2-4).

**Fig. 3:**
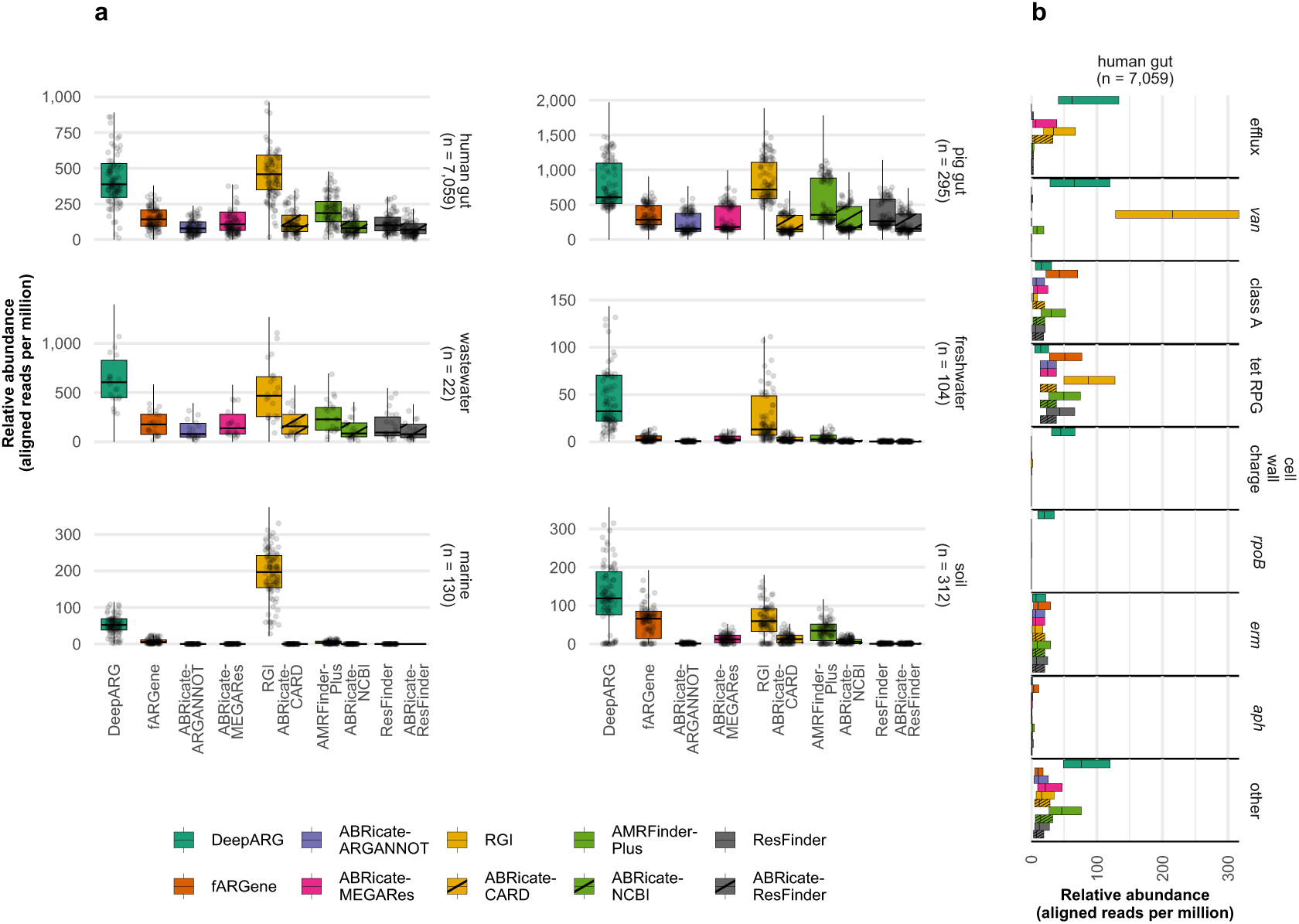
Disparities in ARG relative abundance and functional profiles across detection pipelines. **(a)** Distribution of ARG relative abundance (gene-length–normalized read counts per million mapped reads) by detection pipeline. For illustration, up to 100 randomly subsampled metagenomic samples are plotted per habitat (dots). When comparing ARG richness reported by each pipeline, we observed similar patterns (Extended Data Fig. 5). In the boxplots, the center line denotes the median, the box limits represent the interquartile range (IQR), and the whiskers extend to 1.5 × IQR beyond the first and third quartiles. **(b)** Median relative abundance and interquartile range of ARG classes within human gut metagenomic samples. The groups “class A” and “tet RPG” represent Class A 𝛽-lactamases and tetracycline ribosomal protection genes, respectively. Full gene class names and abbreviations are listed in Supplementary Table 2.

The class-specific relative abundance varied substantially across detection pipelines. In the human gut, DeepARG and RGI estimated median relative abundances for *van* genes and efflux pumps at least seven-fold higher than those of all other pipelines. Furthermore, DeepARG reported a high abundance associated with *rpoB* and cell wall charge genes. Other gene classes, including 𝛽-lactamases, macrolide resistance genes (*erm*), and aminoglycoside phosphotransferases (*aph*), were primarily reported by fARGene and AMRFinderPlus (Fig. 3b; Extended Data Figs. 2-4).

### Variation in the composition of the pan- and core-resistomes

Estimates of the pan-resistome, the collection of ARGs in a habitat, varied substantially across pipelines. The largest pan-resistomes were estimated by DeepARG and RGI, with median sizes 2.5- to 32-fold larger than those of other pipelines across host-associated habitats (Fig. 4a and Extended Data Fig. 6). Consistent with our findings on total ARG counts, the number of genes in the pan-resistome differed even when pipelines used the same underlying reference database (Fig. 4a and Extended Data Fig. 6).

**Fig. 4:**
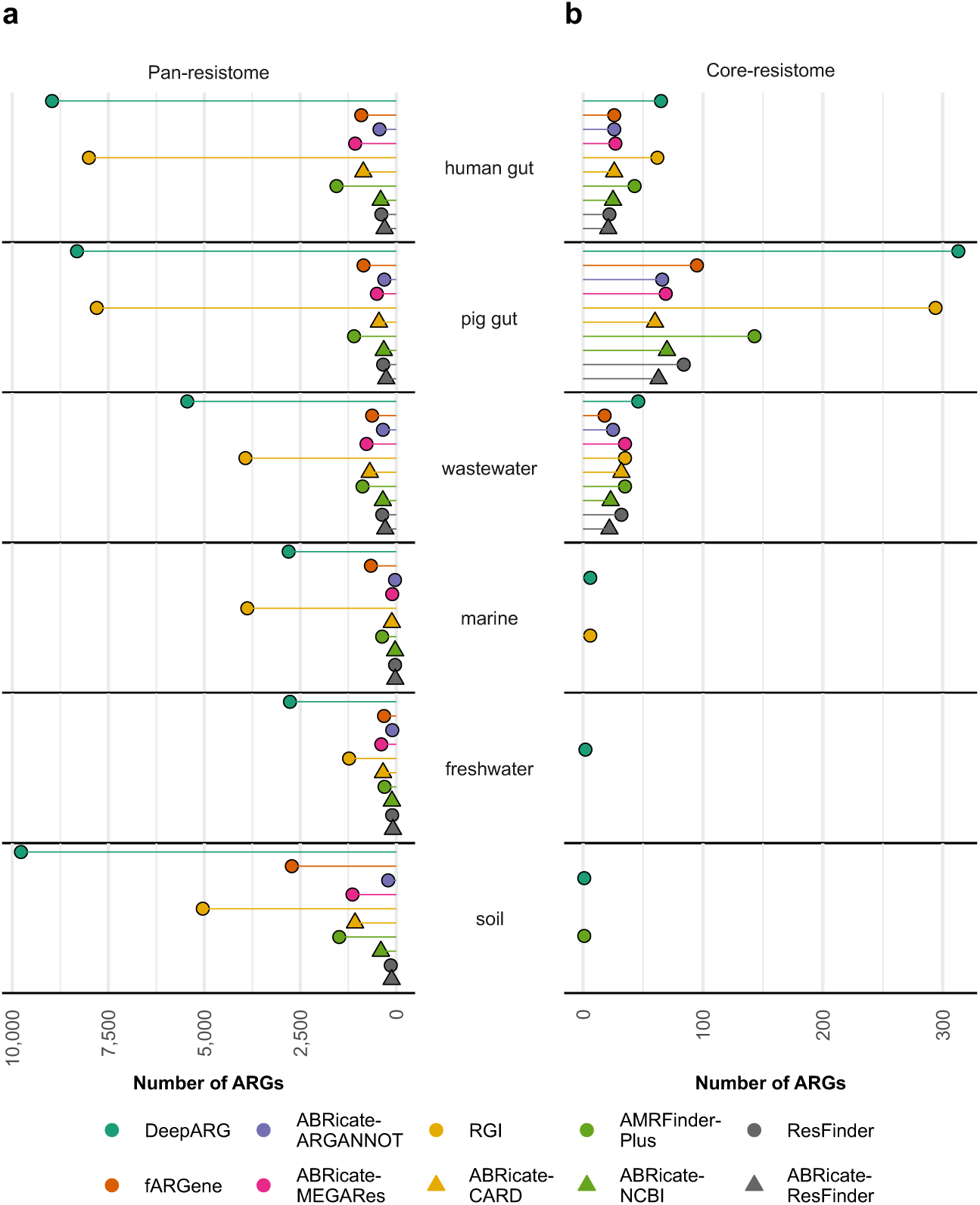
Comparison of pan- and core-resistomes across detection pipelines and habitats. Total number of ARGs reported within the **(a)** pan- and **(b)** core-resistomes for each habitat, stratified by detection pipeline.

A core-resistome, the subset of ARGs commonly present within a habitat, was consistently reported by all pipelines in the mammalian gut, human oral, and wastewater habitats, whereas it was not consistently observed in external habitats (Fig. 4b and Extended Data Fig. 6). In gut habitats, core-resistome estimates from DeepARG and RGI were, on average, 5.3-fold larger than those estimated from other methods. Moreover, estimates of the core-resistome in the marine environment from these two pipelines included six efflux pump genes (two shared between both pipelines), two *rpoB* genes from DeepARG, and two *van* genes from RGI, whereas no core-resistome genes were identified by any other pipeline.

Substantial class-specific differences were also observed in resistome composition across habitats. In the human gut, DeepARG and RGI primarily reported *rpoB* and *van* genes, most of which were reported only by these two pipelines (Extended Data Fig. 7; Supplementary Table 3). Notably, of the 127 ARGs defined as core to the human gut by at least one pipeline, 78 (61%) were identified exclusively by either RGI or DeepARG. We also identified a consensus core-resistome of 24 ARGs reported by at least five pipelines, comprising mainly tetracycline RPGs (10), *erm* genes (5), and Class A 𝛽-lactamases (4). However, even within these shared gene classes, RGI reported twice as many tetracycline RPGs as any other pipeline.

### Cross-Pipeline ARG Class Coverage

To quantify consistency across pipelines and identify systematic differences at the gene-class level, we developed the class-specific coverage (CSC) measure. For a given ARG class, the CSC of pipeline 𝐴 with respect to pipeline 𝐵 is the proportion of ARGs reported by pipeline 𝐵 that were also reported by pipeline 𝐴. While analogous to recall, we intentionally use the term “coverage” to avoid implying that any single pipeline constitutes a ground truth.

fARGene and AMRFinderPlus obtained the highest CSC, with median across gene classes and pipelines of 97% and 88%, respectively (Fig. 5; Extended Data Fig. 8). The higher CSC of these pipelines is likely attributable to their use of Hidden Markov Models (HMMs), which provide enhanced sensitivity in sequence identification compared to alignment^53^.

**Fig. 5:**
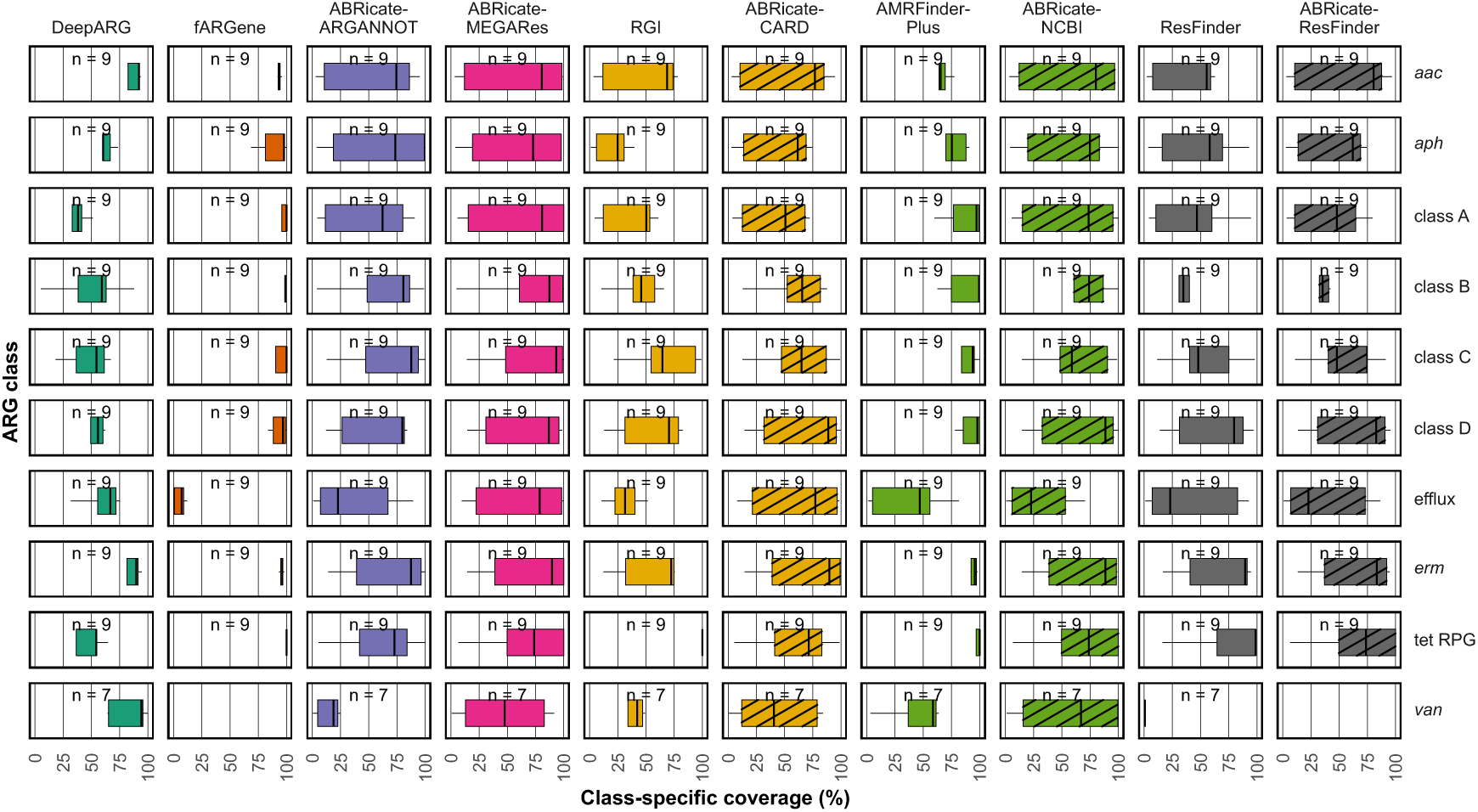
Pipeline class-specific coverage (CSC). Distribution of the CSC by individual ARG functional class and pipeline. In all boxplots, the center line denotes the median, the box limits represent the interquartile range (IQR), and the whiskers extend to 1.5 × IQR beyond the first and third quartiles of the CSC for each pipeline compared with the rest by class. The groups “class A”, “class B”, “class C”, and “class D” represent 𝛽-lactamases, while “tet RPG” represents tetracycline ribosomal protection genes. Full gene class names and abbreviations are listed in Supplementary Table 2.

fARGene exceeded a median of 93% CSC across pipelines for most gene classes except pumps (7%), likely because its efflux pump model is restricted to tetracycline-specific efflux pumps^36^. In contrast, RGI exhibited remarkably low CSC in its two largest classes, *van* and efflux pumps, with median values across pipelines of 42% and 32%, respectively, despite the large number of genes reported by this pipeline belonging to these classes (58,687) and its permissive thresholds (third quartile identity level of 46% relative to the reference database). The comprehensiveness of the MEGARes database was supported by the high CSC of ABRicate-MEGARes relative to other ABRicate-based pipelines (median across pipelines = 100%); however, CSC decreased markedly compared with non-ABRicate pipelines (median 50% across pipelines and gene classes).

## Discussion

Fundamentally, ARG detection pipelines are developed with distinct objectives. While one can define an ARG as a gene whose presence confers resistance^54^, there are multiple valid operationalizations of this concept. Furthermore, reporting ARGs beyond exact sequence matches entails a trade-off between precision and recall, with no optimal answer. Pipelines prioritizing recall over precision may cause undue alarm, whereas prioritizing precision may fail to identify novel *bona fide* ARGs. Cross-study comparisons should take these differences into account.

In this context, DeepARG and RGI can be considered exploratory pipelines that include low-identity homologs, thereby producing larger sets of latent ARGs. In contrast, ResFinder and most ABRicate variants use more stringent reporting criteria, resulting in smaller ARG sets. *A priori*, high-identity results are more likely to be functional, but low identity results from fARGene in environmental data have been shown to be functional^10,32^. We observed, however, that the larger sets reported by DeepARG and RGI do not include a large fraction of the results of the more stringent pipelines, even when the same underlying database was used.

A limitation of this study is that we evaluated a single metagenomic processing strategy: annotating assembled coding sequences with abundances derived by mapping short reads to a non-redundant catalogue^55^. Alternative approaches, such as directly annotating short reads^29,56–58^, would likely introduce additional variability. Moreover, the absence of a representative ground truth dataset for resistomes limits the ability to benchmark pipeline performance. Consequently, we focused on overlaps between outputs. Nonetheless, the observed variation across pipelines was large enough to lead to contrasting biological conclusions.

Given these results, we recommend that no single pipeline be treated as authoritative. Instead, users should consider whether their approach prioritizes exploratory breadth or high-confidence prediction, and report computational predictions as latent rather than definitive findings in the absence of experimental validation. The distinction matters most in habitats such as soil, where latent ARGs substantially outnumber established ones^18,59^. We believe that a constrained methodological pluralism, whereby multiple approaches are seen as valid, but each carries its own limitations that must be respected and communicated with care, is the most honest reflection of the current state of the art.

## Methods

### Data retrieval

We retrieved 278,788,551 nucleotide and protein unigene sequences from the Global Microbial Gene Catalogue (GMGCv1)^44^. These unigenes represent 95%-identity clusters derived from 2.3 billion predicted open reading frames with a greater than zero estimated abundance in 11,538 GMGCv1-metagenomes derived from habitats of interest, namely human-associated habitats (gut, skin, nose, oral cavity, and vagina); animal gut habitats (cat, dog, mouse, and pig); and external habitats (wastewater, marine, freshwater, and soil).

### Computational identification of ARGs

For nucleotide-based ARG identification, the unigenes were processed with RGI v6.0.3^23^ using DIAMOND aligner against the CARD v4.0.0 database; fARGene v0.1^60^, covering all the major classes of beta-lactamases, aminoglycoside resistance, macrolide resistance, quinolone resistance, and tetracycline resistance^31–36^; DeepARG v2^29^; ABRicate v1.0.1 with ARG-ANNOT, CARD, MEGARes v2.0, NCBI, and ResFinder databases (all updated 14 January 2025)^23,27,41,42,61^; and ResFinder v2.4.0^28^. For protein-based ARG identification, the unigenes were analyzed using AMRFinderPlus v4.0.15 with the 2024-12-18.1 database^37^. All tools were run with default parameters. We filtered the ARGs from each tool to include unigenes present in at least one of the metagenomes from the habitats described above (with estimated abundance greater than zero). Overall, 178,107 unigenes were reported as ARG by at least one of these ten pipelines and were detected across the 13 habitats. These pipelines and unigenes constitute the main results. For intra-tool comparison, we processed the unigenes’ nucleotide sequences with RGI^23^ using BLAST aligner and AMRFinderPlus, and the unigenes’ protein sequences with RGI^23^ using DIAMOND aligner, fARGene, and DeepARG. An additional 4,914 unigenes were reported as ARGs.

Predictions based on nucleotide or amino acid sequences were highly consistent across most tools. For DeepARG, fARGene, and RGI-DIAMOND, more than 99.5% of the ARGs identified from nucleotide sequences were also detected from amino acid sequences (Extended Data Fig. 9a). In contrast, AMRFinderPlus, using nucleotide sequences, detected only 14% of the ARGs this tool reported using amino acid sequences (by the tool’s design, none of the AMRFinderPlus HMM models were applied to nucleotide sequences, Extended Data Fig. 9a). Similarly, when run on nucleotide sequences, RGI using DIAMOND or BLAST reported only 91% of the total ARGs detected by this tool across all parameter settings (Extended Data Figs. 1a). ABRicate shows low concordance to AMRFinder and NCBI, but a high overlap to ResFinder (Extended Data Fig. 9b). ABRicate-MEGARes showed higher concordance, recovering up to 88% of ARGs from other ABRicate-based runs (Extended Data Figs. 9c and 9d).

### Ontology harmonization

ARG detection results were harmonized to allow cross-pipeline comparison. The outputs of DeepARG, AMRFinderPlus, ABRicate, and ResFinder were processed with argNorm v1.0.0^21^. Missing ARO identifiers in ABRicate-CARD outputs were assigned by mapping gene names to CARD model names (v4.0.0) or, if still absent, manually via the Ontology Lookup Service^40^. Genes from ResFinder or ABRicate (ResFinder, MEGARes v2.0, NCBI datasets) that lacked AROs were assigned via RGI with DIAMOND using loose matching. fARGene ARGs were manually annotated based on the model that flagged the gene. Identity levels for fARGene predictions were obtained via RGI alignments with DIAMOND, providing a cross-check of automatic assignments. All ARGs with AROs were further annotated with two hierarchical levels: (1) a parent level from the Ontology Lookup Service to consolidate genes into broader classes, and (2) a manually curated higher level aggregating parent classes into unified categories (Supplementary Table 2). We further merged the Major Facilitator Superfamily (MFS) efflux pump class with all other efflux pump classes.

### Concordance analysis

The pairwise concordance was calculated using the Jaccard index for ARG prediction sets across all pipeline pairs. The Jaccard index was calculated as 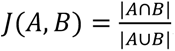, for the set of ARGs, 𝐴 and 𝐵, reported by any pipelines A and B.

Let 𝐴 be the set of unigenes reported by pipeline A, and 𝐴_&_the subset of those annotated with gene class 𝑔. We defined the class-specific coverage (CSC) of pipeline A for gene class 𝑔 with respect to pipeline B as 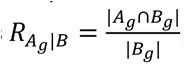, where 𝐵*_g_* is the set of unigenes reported by pipeline 𝐵 and belonging to gene class 𝑔. For the edge case where a unigene is reported by both pipelines but assigned to different ARG classes (this happens for 377 unigenes; 0.21% of the total reported ARGs), CSC calculations were performed by harmonizing class assignments to those of pipeline 𝐴 in the pairwise comparison. This was particularly relevant for aminoglycoside resistance genes, some of which encode bifunctional enzymes and can therefore receive different but valid class assignments depending on the reference gene, as well as for genes classified as both efflux pumps and other functional variants. If either |𝐴_&_| = 0 or |𝐵_&_| = 0, the CSC was not computed.

### Quantification of the abundance and richness of ARGs

We retrieved unigene abundances from GMGCv1, using the normed10m estimate, which includes normalisation for both unigene length and total number of read counts per sample, distributing multiple-mappers according to the distribution of unique mappers^62^. The richness was calculated as the number of unique ARGs identified in each metagenomic sample after rarefaction to 5,000,000 reads. Detailed methods for abundance estimation are described in Coelho et al.^44^.

### Estimation of the pan- and core-resistomes

Following the definitions of the pan- and core-resistomes from Inda-Díaz et al.^18^, genes were clustered at 90% nucleotide identity using VSEARCH v2.30.0^63^, and cluster centroids were used for downstream analyses. For each pipeline and habitat, the core-resistome was estimated by randomly selecting 500 subsamples of 100 metagenomic samples. For each subsample, we recorded (1) the subsample core, i.e., the gene class of the centroids with a detection value ≥1 in at least 50% of samples, grouped by habitat, and (2) the total number of centroids by ontology with a detection value ≥1 in any sample, as a measure of richness. The pan-resistome size for each habitat and ontology level was calculated as the mean richness across the 500 subsamples. The core-resistome corresponded to ARGs present in ≥90% of the subsample cores. Habitats with fewer than 100 samples included all samples in the computation. The pan- and core-resistomes were calculated after rarefaction to 5,000,000 reads per metagenomic sample. Results using alternative thresholds for core-resistome definition are provided on the *ARG-pipelines* website (https://arg-pipelines.big-data-biology.org).

## Data availability

The output of the ARG detection pipelines and normalization, as well as the estimated abundance, richness, and sample metadata, have been deposited at Zenodo (https://doi.org/10.5281/zenodo.19702877). The source unigene sequences, metagenomic sample annotations, and abundances are available on the Global Microbial Gene Catalogue website (https://gmgc.embl.de/). An interactive visualization website of the results is available on the *ARG-pipelines* website (https://arg-pipelines.big-data-biology.org).

## Code availability

The code used to run the ARG-detection pipelines, to generate the figures, and for the supporting website are available in the GitHub repository (https://github.com/BigDataBiology/IndaDiaz2026 ARGTools)

## Acknowledgements

We thank Andy Leu and Catarina Sales e Santos Loureiro from the Queensland University of Technology, and Willem van Schaik from the University of Birmingham for helpful comments on previous versions of this manuscript. We also thank members of the Coelho and Woodcroft groups at the Queensland University of Technology, as well as the EMBARK and SEARCHER consortia, for their helpful comments during the project’s development.

The icons *excess-weight-male-clothed*, *mouse-gray*, *pig-white*, and *dog* in the abstract figure were adapted from Servier Medical Art (https://smart.servier.com) and are licensed under CC BY 3.0 Unported (https://creativecommons.org/licenses/by/3.0/).

## Contributions

J.I., S.U.P., and L.P.C. conceptualized the article and designed the study. J.I., F.A., U.L., V.J., Y.D., J.B.P., and S.U.P. performed computational analyses. J.I. and F.A. visualised the data.

J.I. drafted the first version of the manuscript. All authors edited and approved the final manuscript.

## Extended Data

**Extended Data Fig. 1.**
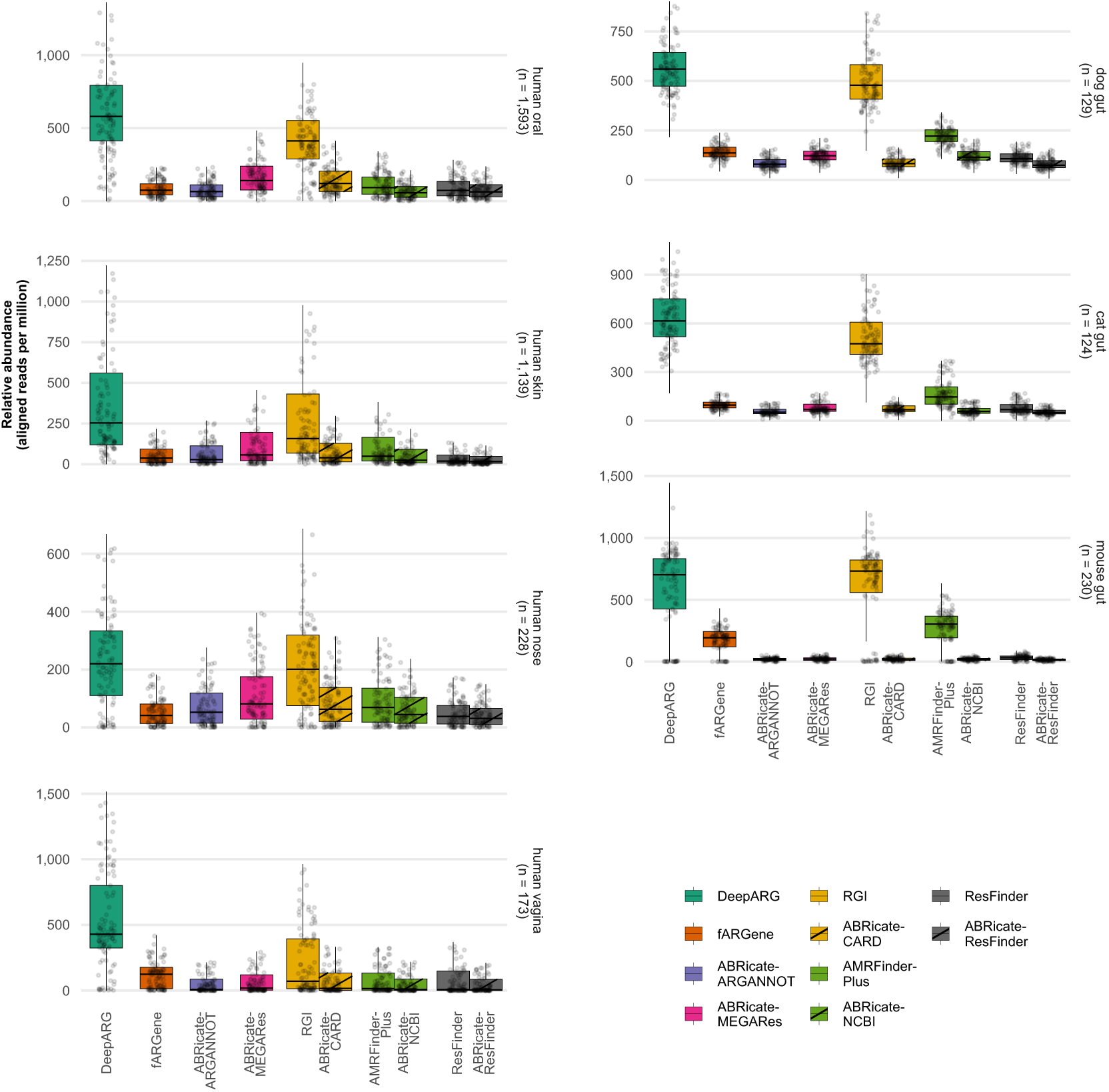
Disparities in ARG relative abundance. Distribution of ARG relative abundance (gene-length–normalized read counts per million mapped reads) as estimated by each detection pipeline. For illustration, up to 100 randomly subsampled metagenomic samples are plotted per habitat (dots). In the boxplots, the center line denotes the median, the box limits represent the IQR, and the whiskers extend to 1.5 × IQR beyond the first and third quartiles.

**Extended Data Fig. 2.**
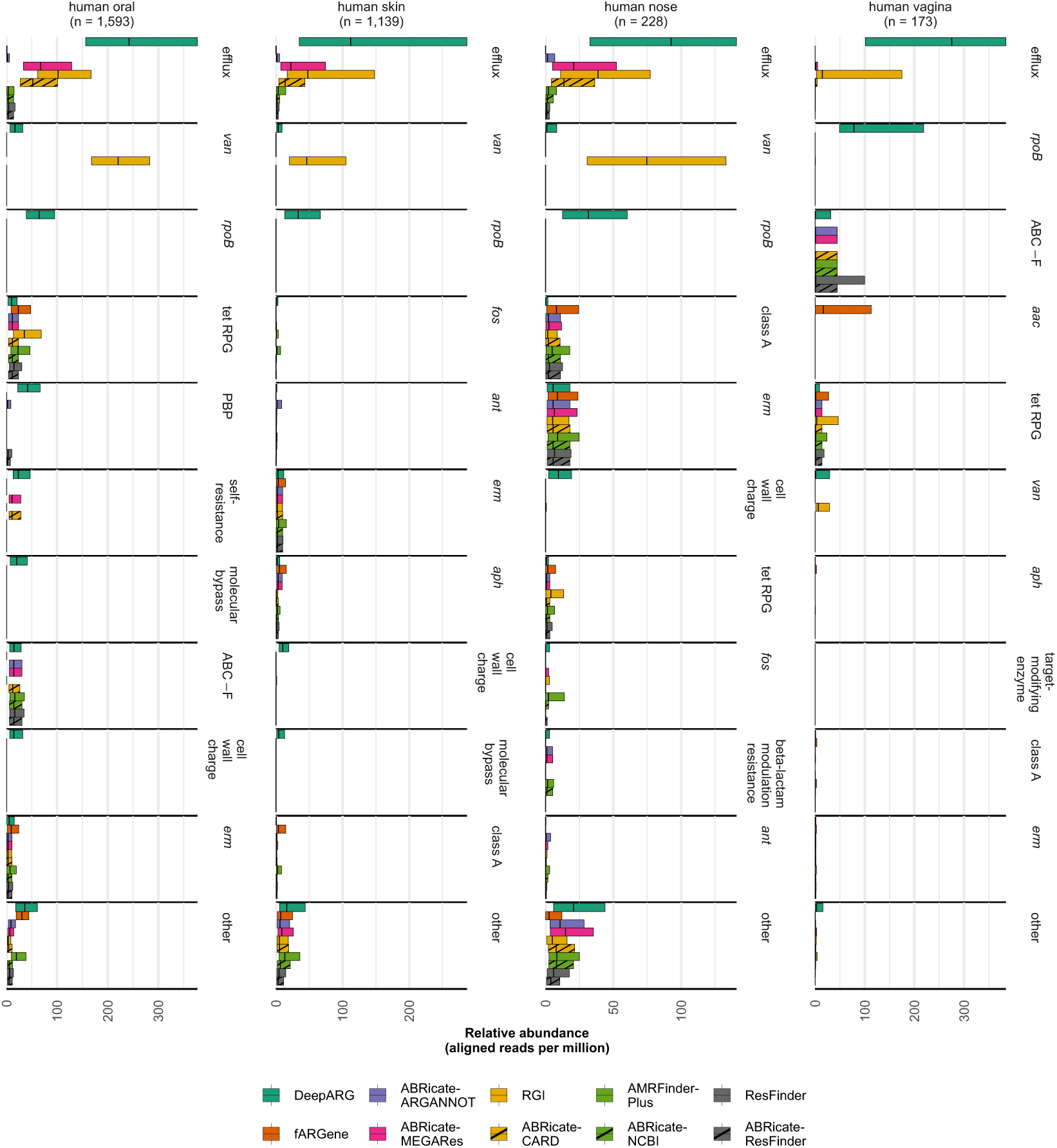
Disparities in ARG relative abundance across functional profiles, detection pipelines, and human microbiomes. Median relative abundance (gene-length–normalized read counts per million mapped reads) and interquartile range of ARG classes within human microbiomes. The groups “class A” and “tet RPG” represent 𝛽-lactamases class A and tetracycline ribosomal protection genes, respectively. Full gene class names and abbreviations are listed in Supplementary Table 2.

**Extended Data Fig. 3.**
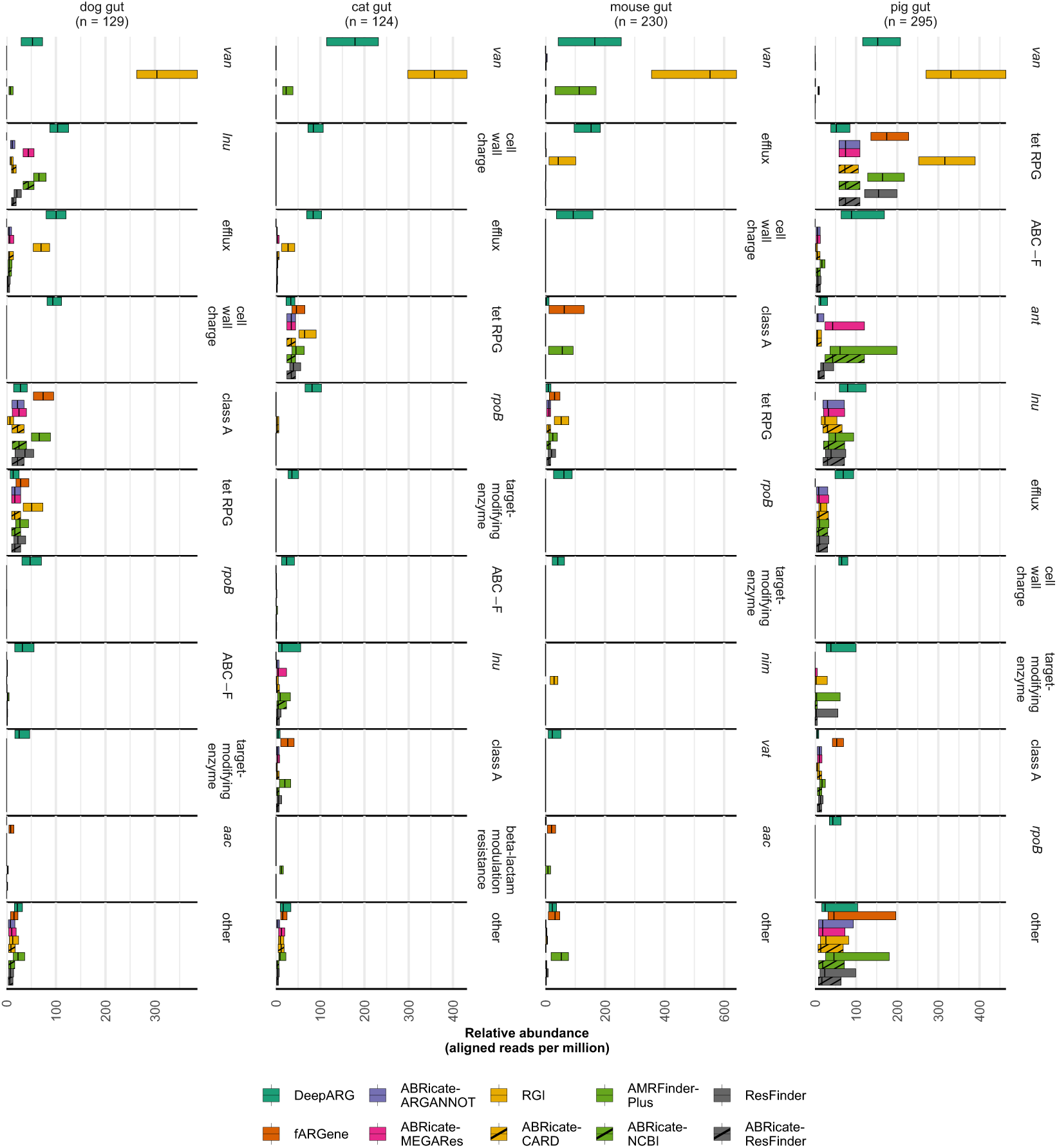
Disparities in ARG relative abundance across functional profiles, detection pipelines, and animal gut microbiomes. Median relative abundance (gene-length–normalized read counts per million mapped reads) and interquartile range of ARG classes within animal gut microbiomes. The groups “class A” and “tet RPG” represent 𝛽-lactamases class A and tetracycline ribosomal protection genes, respectively. Full gene class names and abbreviations are listed in Supplementary Table 2.

**Extended Data Fig. 4.**
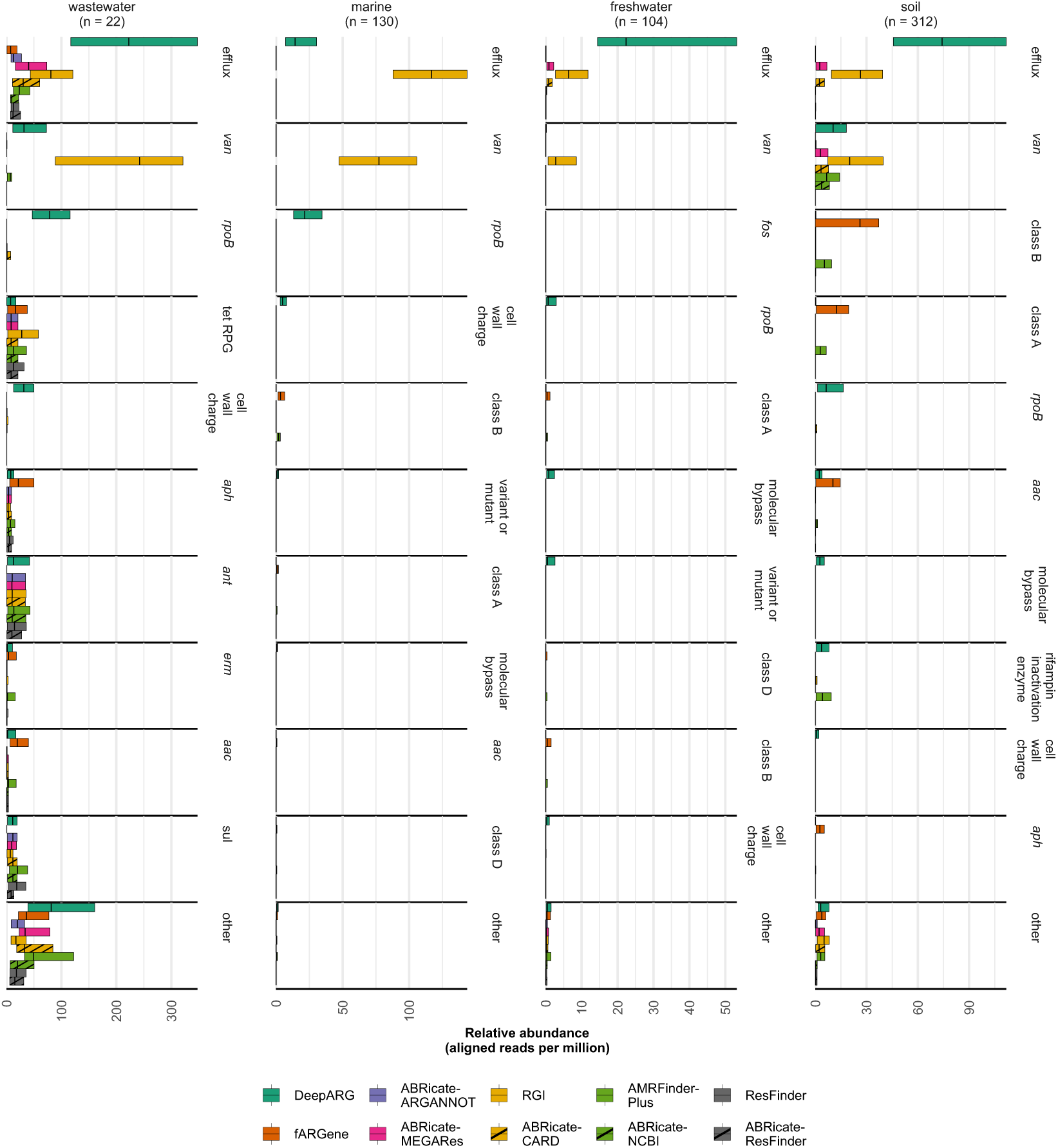
Disparities in ARG relative abundance across functional profiles, detection pipelines, and the microbiomes of external environments. Median relative abundance (gene-length–normalized read counts per million mapped reads) and interquartile range of ARG classes within the microbiomes of external environments. The groups “class A” and “tet RPG” represent 𝛽-lactamases class A and tetracycline ribosomal protection genes, respectively. Full gene class names and abbreviations are listed in Supplementary Table 2.

**Extended Data Fig. 5.**
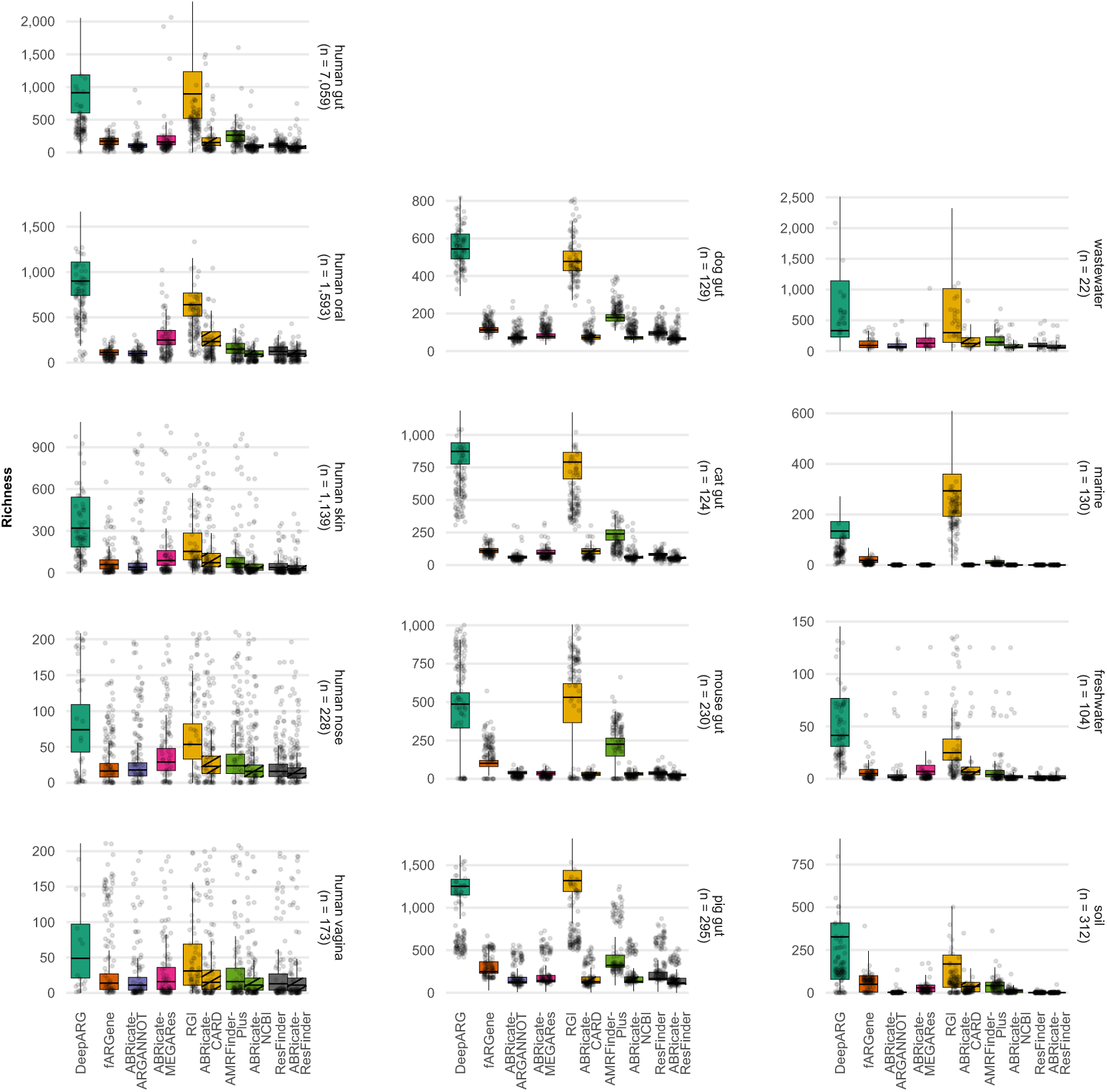
Disparities in ARG richness across detection pipelines and habitats. Distribution of the richness of ARG, the number of unique ARGs identified within a metagenomic sample, as estimated by each detection pipeline. For illustration, up to 100 randomly subsampled metagenomic samples are plotted per habitat (dots). In the boxplots, the center line denotes the median, the box limits represent the IQR, and the whiskers extend to 1.5 × IQR beyond the first and third quartiles.

**Extended Data Fig. 6.**
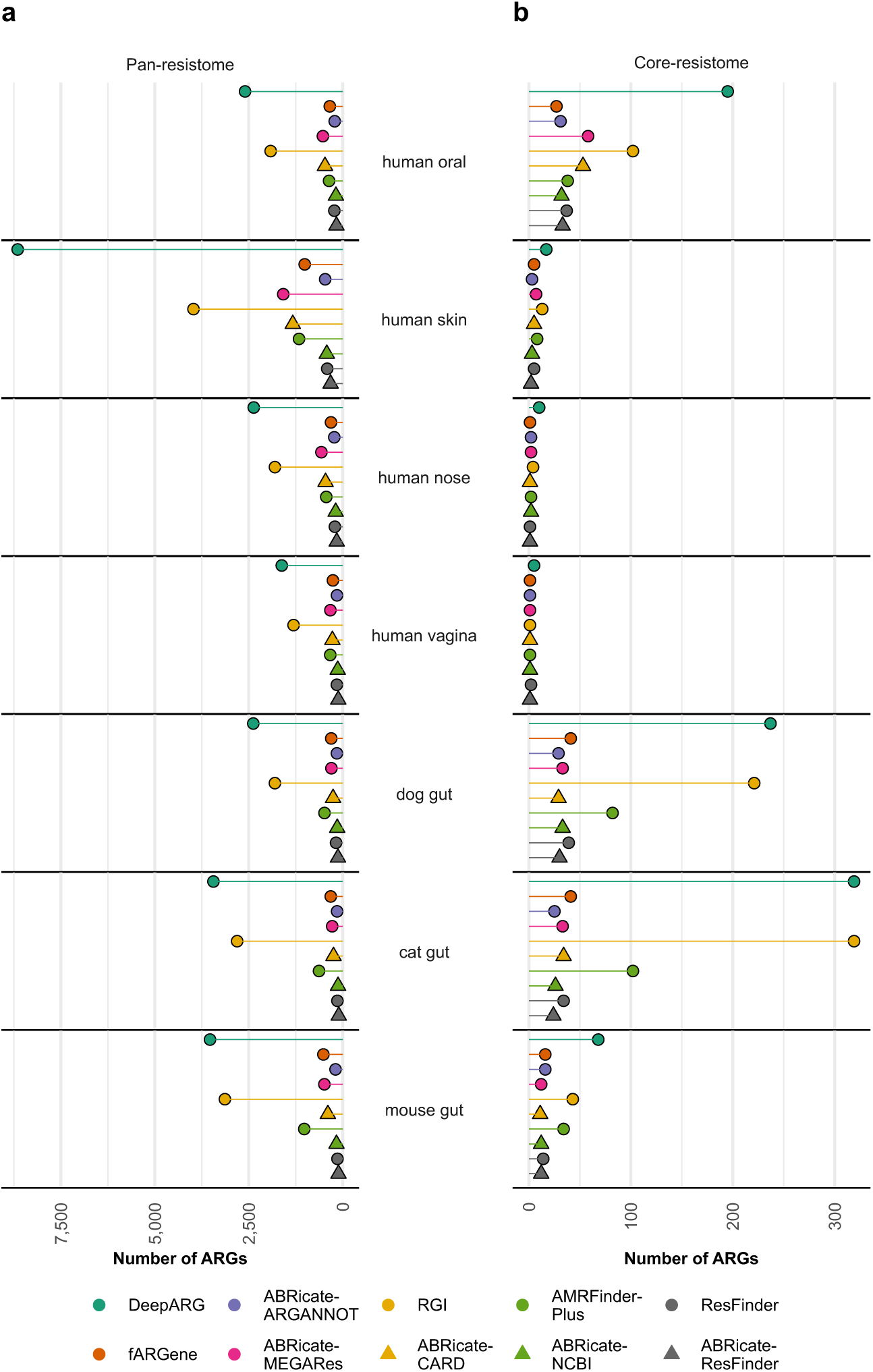
Comparison of pan- and core-resistomes across detection pipelines and habitats. Total number of ARGs reported within the **(a)** pan- and **(b)** core-resistomes for each habitat, stratified by detection pipeline.

**Extended Data Fig. 7.**
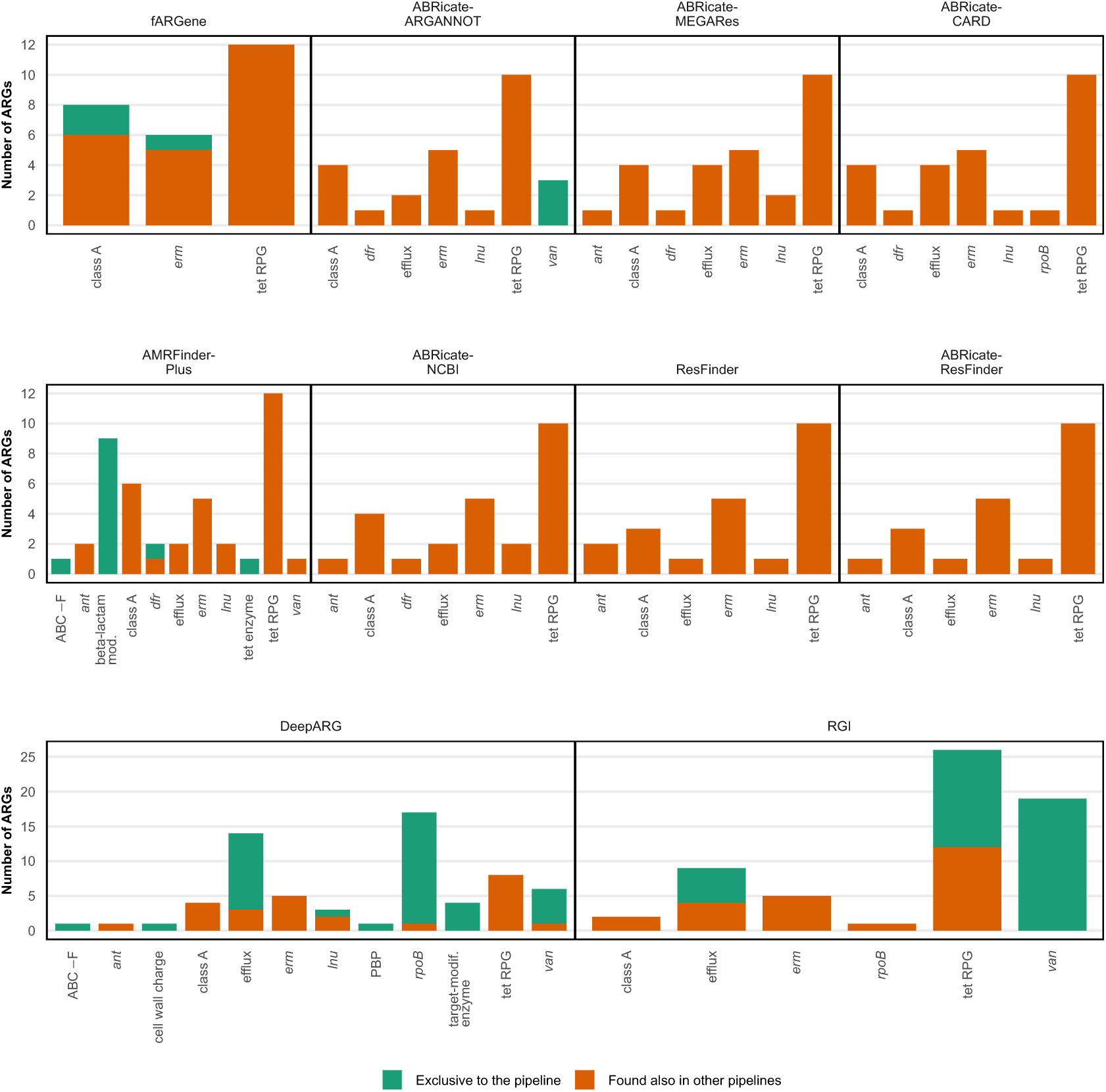
Composition and overlap of the human core-resistome. Estimated number of genes in the core-resistome by gene class, pipeline, and if the ARG was found in the core-resistome estimated by one exclusive pipeline or more. The groups “class A”, “class B”, “class C”, and “class D” represent 𝛽-lactamases, while “tet RPG” represents tetracycline ribosomal protection genes. Full gene class names and abbreviations are listed in Supplementary Table 2.

**Extended Data Fig. 8:**
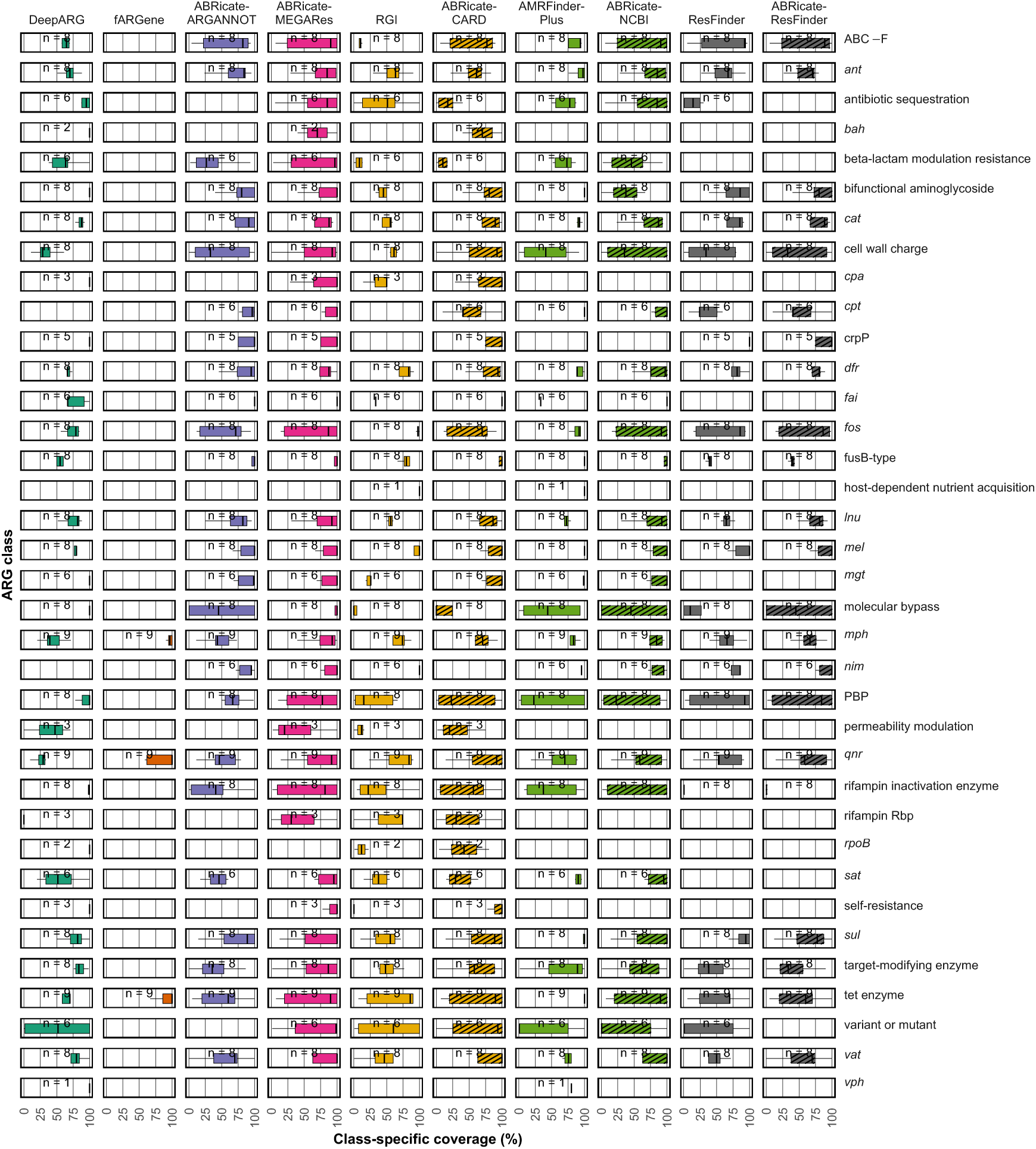
Pipeline concordance as class-specific coverage (CSC) of ARG predictions. Breakdown of CSC by individual ARG functional class (y-axis), illustrating the varying relative sensitivity of each pipeline (x-axis) with respect to other pipelines. In all boxplots, the center line denotes the median, the box limits represent the interquartile range (IQR), and the whiskers extend to 1.5 × IQR beyond the first and third quartiles. The groups “class A”, “class B”, “class C”, and “class D” represent 𝛽-lactamase subclasses, while “tet RPG” represents tetracycline ribosomal protection genes. Full gene class names and abbreviations are listed in Supplementary Table 2.

**Extended Data Fig. 9.**
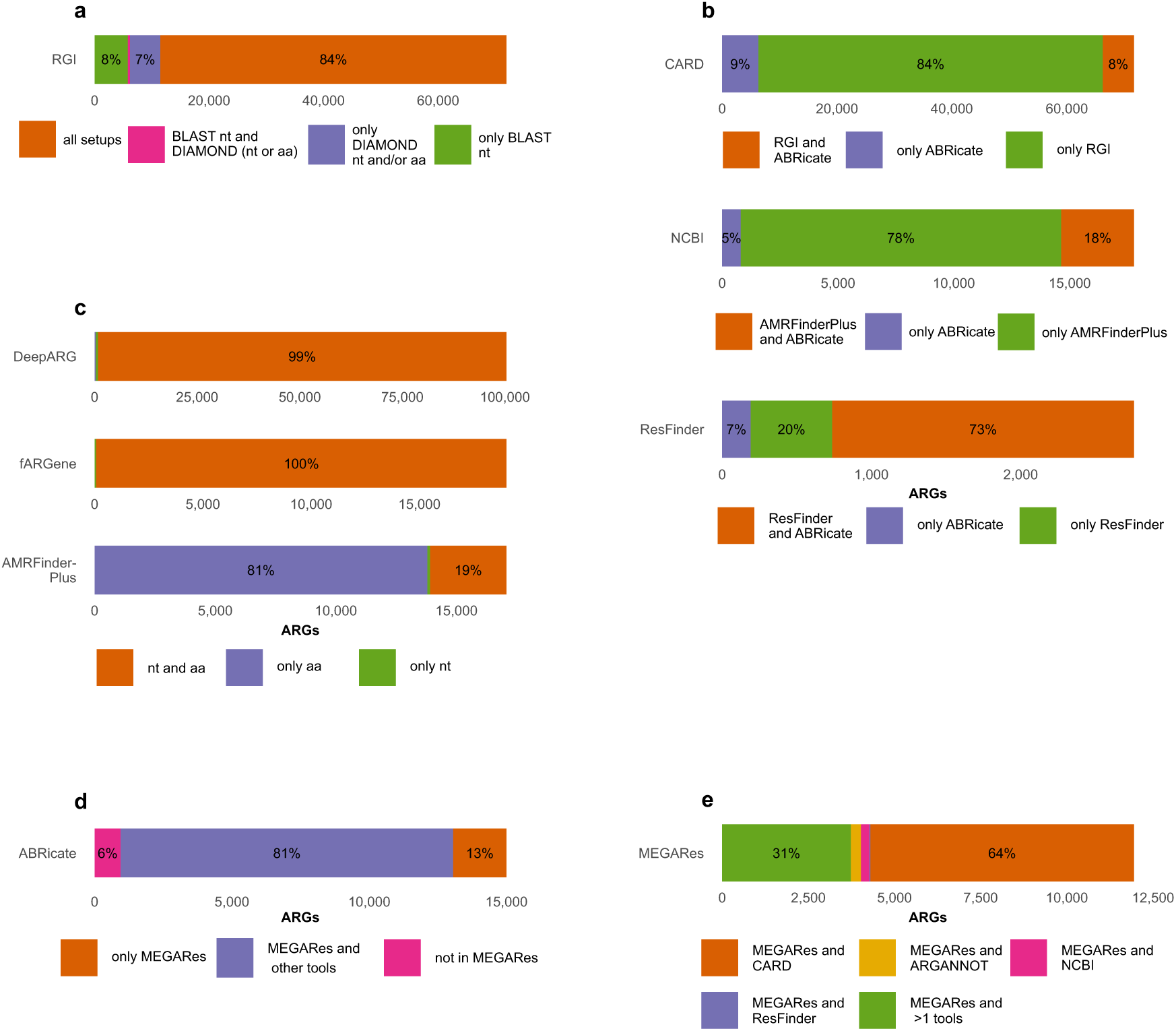
Prediction overlap. Intra-tool overlap for **(a)** RGI, **(c)** DeepARG, fARGene, and AMRFinderPlus. The number of ARGs reported exclusively by each sequence format and/or aligner combination per tool, and the overlap among them. The sequence format is specified as nucleotide (nt) or amino acid (aa). **(b)** Inter-tool overlap for tools using similar reference databases. The number of ARGs reported exclusively by each tool and their overlap across ARG databases. RGI with DIAMOND aligner and ResFinder in using nucleotide sequences, and AMRFinderPlus using amino acid sequences, are compared to ABRicate using nucleotide sequences and the respective databases. **(d)** Intra-tool ARGs overlap in ABRicate. The number of ARGs reported exclusively by ABRicate with the specified database versus MEGARes. **(e)** Overlap between MEGARes and other databases used by ABRicate.

## Supplementary information

Supplementary Table 1. Metagenomic samples per habitat used in this study (file TableS1.csv).

Supplementary Table 2. Aggregation of genes from ARO to the ARG classes used here (file TableS2.csv).

Supplementary Table 3. Human gut core-resistome unigenes, annotated by ARG class and detection pipeline (file TableS3.csv).

All figures and supplementary materials can be fully accessed and generated through the interactive visualization website *ARG-pipelines* (https://arg-pipelines.big-data-biology.org).

